# PTEN inhibition promotes robust growth of bulbospinal respiratory axons and partial recovery of diaphragm function in a chronic model of cervical contusion spinal cord injury

**DOI:** 10.1101/2024.01.10.575021

**Authors:** Pauline Michel-Flutot, Lan Cheng, Samantha J Thomas, Brianna Lisi, Harrison Schwartz, Sandy Lam, Megan Lyttle, David A Jaffe, George Smith, Shuxin Li, Megan C Wright, Angelo C Lepore

**Author notes:** Co-first authors: these authors contributed equally to this work (Pauline Michel-Flutot, Lan Cheng). Corresponding author: Angelo C. Lepore, Ph.D., Department of Neuroscience, Vickie and Jack Farber Institute for Neuroscience, Sidney Kimmel Medical College at Thomas Jefferson University, 233 S. 10th St., Bluemle Life Sciences Building Room 245, Philadelphia, PA 19107.

## Abstract

High spinal cord injury (SCI) leads to persistent and debilitating compromise in respiratory function. Cervical SCI not only causes the death of phrenic motor neurons (PhMNs) that innervate the diaphragm, but also damages descending respiratory pathways originating in the rostral ventral respiratory group (rVRG) located in the brainstem, resulting in denervation and consequent silencing of spared PhMNs located caudal to injury. It is imperative to determine whether interventions targeting rVRG axon growth and respiratory neural circuit reconnection are efficacious in chronic cervical contusion SCI, given that the vast majority of individuals are chronically-injured and most cases of SCI involve contusion-type damage to the cervical region. We therefore employed a clinically-relevant rat model of chronic cervical hemicontusion to test therapeutic manipulations aimed at reconstructing damaged rVRG-PhMN-diaphragm circuitry to achieve recovery of respiratory function. At a chronic time point post-injury, we systemically administered: an antagonist peptide directed against phosphatase and tensin homolog (PTEN), a central inhibitor of neuron-intrinsic axon growth potential; an antagonist peptide directed against receptor-type protein tyrosine phosphatase sigma (PTPσ), another important negative regulator of axon growth capacity; or a combination of these two peptides. PTEN antagonist peptide (PAP4) promoted partial recovery of diaphragm motor activity out to nine months post-injury, while PTPσ peptide did not impact diaphragm function after cervical SCI. Furthermore, PAP4 promoted robust growth of descending bulbospinal rVRG axons caudal to the injury within the denervated portion of the PhMN pool, while PTPσ peptide did not affect rVRG axon growth at this location that is critical to control of diaphragmatic respiratory function. In conclusion, we find that, when PTEN inhibition is targeted at a chronic time point following cervical contusion that is most relevant to the SCI clinical population, our non-invasive PAP4 strategy can successfully promote significant regrowth of damaged respiratory neural circuitry and also partial recovery of diaphragm motor function.

**HIGHLIGHTS:**

- PTEN antagonist peptide promotes partial diaphragm function recovery in chronic cervical contusion SCI.
- PTPσ inhibitory peptide does not impact diaphragm function recovery in chronic cervical contusion SCI.
- PTEN antagonist peptide promotes growth of bulbospinal rVRG axons in chronic cervical contusion SCI.
- PTPσ peptide does not affect rVRG axon growth in chronic cervical contusion SCI.

## INTRODUCTION

Spinal cord injury (SCI) leads to permanent motor, sensory and autonomic dysfunction (Rowland et al 2008, Venkatesh et al 2019). Furthermore, SCI occurs most frequently at the cervical level, which often results in long-lasting respiratory impairment (Ahuja et al 2017, Winslow & Rozovsky 2003). To date, a number of therapeutic approaches have shown beneficial effects for stimulating functional recovery when delivered at early time points following injury (Thuret et al 2006). However, there has been very limited success with developing effective treatments to achieve significant spinal cord repair in the context of chronic cervical SCI.

Several important mechanisms have been identified that limit regrowth and reconnection of damaged circuits in the adult central nervous system (CNS). Among them, neuronal-intrinsic inhibition of axon growth capacity plays a particularly important role (Luo & Park 2012). In addition, neuronal-extrinsic factors also limit axon growth due to modification of extracellular network composition after SCI (Giger et al 2010).

Phosphatase and tensin homolog (PTEN) is a neuronal-intrinsic inhibitor of the phosphatidylinositol 3-kinase (PI3K)/Akt/mammalian target of rapamycin (mTOR) pathway that is involved in regulation of axon growth ability (Park et al 2008). A number of studies have shown that inhibiting PTEN allows spared and injured axons to extend following CNS injury, leading to some functional recovery in various SCI animal models (Bhowmick & Abdul-Muneer 2020, Cheng et al 2021a, Cho et al 2020, Danilov & Steward 2015, Du et al 2015, Lewandowski & Steward 2014, Liu et al 2012, Liu et al 2010, Metcalfe & Steward 2023, Ohtake et al 2014, Stewart et al 2023, Urban et al 2019, Walker & Xu 2014, Zhu et al 2017, Zukor et al 2013). We previously developed a novel PTEN inhibitory peptide (*i.e.*, PAP4 for “PTEN antagonist peptide 4”) that can cross the blood brain barrier when administered systemically (Ohtake et al 2014). PAP4 is a 30-amino acid peptide that binds to the C-terminal portion of PTEN, resulting in PTEN inhibition. We systemically administered PAP4 in a rat model of C2 spinal cord hemisection (C2Hx) and evaluated its effects on the neurorespiratory system (Cheng et al 2021a, Urban et al 2019). We showed that, when delivered sub-acutely following C2Hx, daily PAP4 administration leads to improved diaphragm activity. This is associated with long-distance regeneration of ipsilesional rVRG axons through the lesion and towards denervated PhMNs located distal to the injury site (Urban et al 2019). In view of these encouraging results, we then aimed to determine whether PAP4, when instead administered at a chronic stage following cervical SCI, could also have beneficial effects on respiratory function, which is highly important given that the vast majority of individuals living with SCI are at a chronic stage. We showed that, while chronic PAP4 administration induces robust rVRG axon regeneration in the chronic C2Hx paradigm, only modest diaphragm functional recovery was achieved, which was possibly due to the lack of rVRG axon-PhMN synaptic reconnection that we find occurs in the chronic SCI model but not the sub-acute delivery paradigm (Cheng et al 2021a).

Chondroitin sulfate proteoglycans (CSPGs) are physiologically expressed in the spinal cord extracellular matrix. They limit axon growth, possibly to prevent inappropriate neuronal plasticity in the adult nervous system (Bradbury & Burnside 2019). Following CNS injury, increased CSPG expression occurs in a number of cell types in the spinal cord, including reactive astrocytes and fibroblasts (Bradbury & Burnside 2019). CSPGs signal through receptors present on the axon membrane: protein tyrosine phosphatase sigma (PTPσ) (Shen et al 2009), leukocyte common antigen-related phosphatase (LAR) (Fisher et al 2011), and Nogo receptors 1 and 3 (Dickendesher et al 2012). The interaction between CSPGs and their receptors leads to intracellular signaling and axon growth inhibition, thereby limiting axon plasticity and functional recovery following SCI (Bradbury & Burnside 2019). Direct removal of these CSPGs by using the bacterial enzyme chondroitinase ABC (ChABC) that selectively cleaves the glycosaminoglycan (GAG) side chains from CSPGs promotes axon sprouting and some functional recovery in preclinical models of high SCI (Alilain et al 2011, Bradbury et al 2002, García-Alías et al 2009, James et al 2015, Ramer et al 2014, Rosenzweig et al 2019, Tester & Howland 2008, Warren et al 2018). Targeting of CSPG receptors has also shown beneficial effects on functional recovery after SCI through therapeutic modulation of PTPσ (Ito et al 2021, Lang et al 2015) or LAR (Cheng et al 2021b, Xu et al 2015) activity/expression. These two receptors belong to the type IIa/LAR subfamily of receptor protein tyrosine phosphatases (RPTP) (Tonks 2006) and bind CSPGs through their extracellular Ig-like domain (Xie et al 2020). We previously tested the effect on respiratory recovery of acute administration of a PTPσ inhibitory peptide in a rat model of C2Hx (administered from day-0 to day-14 post-injury). We demonstrated that this paradigm leads to improved ipsilesional diaphragm activity, which was associated with axon sprouting from spared bulbospinal respiratory neurons located in the contralesional rVRG (Urban et al 2020). While this PTPσ peptide administration approach showed beneficial effects when delivered during the acute and sub-acute period, it remains to be determined whether it can exert similar effects when delivered in a chronic SCI setting.

To date, the vast majority of preclinical animal studies have focused on improving functional recovery after SCI by applying therapeutics in acute or sub-acute paradigms. Paradoxically, most affected individuals are living with a chronic SCI. Furthermore, as we have previously shown, therapeutics delivered at early time points post-injury may not produce similarly robust beneficial effects when delivered chronically after SCI (Cheng et al 2021a, Urban et al 2019). This makes the rigorous testing of potential therapeutics in preclinical chronic SCI models an incredibly important and necessary step toward the development of novel treatment strategies.

In this study, we evaluated the functional and neuroanatomical effects of inhibiting both neuronal-intrinsic and neuronal-extrinsic factors that limit axon growth capacity. At a chronic stage following cervical contusion SCI, we systemically delivered PAP4, a PTPσ peptide inhibitor, or a combination of these two peptides. Importantly, we chose to use a cervical contusion SCI over the C2Hx paradigm in this work as this model best represents the type of spinal cord trauma occurring in the clinical population.

## MATERIAL AND METHODS

### Animals

Female Sprague Dawley rats (ENVIGO, USA, n = 51, 250-300 g) were used for this study. All experimental procedures were approved by Thomas Jefferson University Institutional Animal Care and Use Committee (IACUC) committee and conducted in compliance with the National Institutes of Health (NIH) Guide for the Care and Use of Laboratory Animals and the ARRIVE (Animal Research: Reporting of In Vivo Experiments) guidelines. Animals were housed three animals per cage in a temperature, humidity, and light-controlled environment. Food and water were provided *ad libitum* on a 12-hour light/dark cycle. Female animals were used in this study, as we extensively optimized and comprehensively characterized the unilateral cervical contusion SCI model both functionally and histologically in female rats in our previous work (Nicaise et al 2013, Nicaise et al 2012).

### Mid-cervical hemicontusion surgery

All animals received a mid-cervical C4/5 hemicontusion on the right side. Animals were anesthetized with xylazine (10 mg/kg, s.q.; Akorn Inc., Lake Forest, Illinois) and ketamine (120 mg/kg, i.p.; Vedco, Saint Joseph, Missouri). Animals were placed on a heating pad. Briefly, skin and muscle were retracted, then laminectomy and durotomy were performed at the C3 to C5 level (Lepore 2011). Rats then underwent a contusion using the Infinite Horizon Spinal Impactor (Precision Systems and Instrumentation) with a 1.5 mm tip at a force of 395 kdyn (Li et al 2014). Muscles were sutured with 4–0 Vicryl, and wound clips (Braintree Scientific, Braintree, Massachusetts) were used to close the skin. Rats were injected with antisedan (1.2 mg/kg, s.q.) (Zoetis, Parsippany-Troy Hills, New Jersey), lactated Ringer’s solution (10 ml, s. q.) and buprenorphine (0.01 mg/kg, s.q.) (Patterson Veterinary, Greeley, Colorado).

### Rostral ventral respiratory group (rVRG) injection

A subset of animals was microinjected with AAV2-mCherry anterograde tracer in the rVRG ipsilateral to SCI. Animal anesthesia was conducted with injection of ketamine HCl (95.0 mg/kg; i.p.; Vedco, Saint Joseph, MO, USA), and xylazine (10.0 mg/kg; s.c.; Lloyd Laboratories, Shenandoah, IA, USA). C1 vertebra and caudal portion of occipital bone were surgically removed, revealing the obex in the medulla. Animals were then moved to a stereotaxic frame (Kopf Instruments, Tujunga, CA, USA). An UltraMicroPump (World Precision Instruments, Sarasota, FL, USA) injection system with a microsyringe (Hamilton, Reno, NV, USA) attached to a Micro4 Microsyringe Pump Controller (World Precision Instruments, Sarasota, FL, USA) was used to microinject 0.3 ul of AAV2-mCherry (titer: 1.66 × 10^13^ vg/ml). Injection was performed using the following coordinates: 2.0 mm lateral, 1.0 mm rostral, and 2.6 mm ventral to obex (Cheng et al 2021b). After injection completion, the needle was kept in place for 3 min before being removed.

### Compound muscle action potential (CMAP) recordings

Anesthesia was induced with 5% isoflurane (Piramal Healthcare, Bethlehem, Pennsylvania, USA) in 100% oxygen, and maintained through a nose cone with 2–2.5% isoflurane. Animals were placed on a heating pad in supine position. Stimulating electrodes were placed ipsilateral to SCI to stimulate the phrenic nerve, and diaphragm CMAPs were recorded, as previously described (Lepore et al 2010, Martin et al 2015). A single supramaximal stimulus with 0.5 milliseconds duration was delivered using an ADI Powerlab 8/30 stimulator (AD Instruments, Dunedin, New Zealand). At least ten repetitions were performed to obtain the average response, with the recording and stimulating electrodes in the same positions. CMAP recordings were obtained with a BioAMPamplifier (AD Instruments, Dunedin, New Zealand; RRID:SCR_018833), and the amplitude was measured from baseline to peak.

### Retrograde labeling of phrenic motor neurons (PhMNs)

Three days before euthanasia, rats were anesthetized with isoflurane (5% for induction; 2-2.5% for maintenance), and 15–20 μL cholera toxin B subunit (CTB, #104 List Biological Labs, Campbell, California, USA) was bilaterally injected into the intrapleural space to retrogradely label PhMNs, as previously described (Cheng et al 2021a).

### Diaphragm electromyography (EMGdia) recordings

When animals were assessed for diaphragm activity, one of two different types of anesthesia was used: isoflurane or ketamine/xylazine. When isoflurane was used, anesthesia was induced using isoflurane (5% in 21% O2 balanced) in an anesthesia chamber and maintained through a nose cone (2.5% in 100% air balanced). For ketamine/xylazine, animals were anesthetized using intraperitoneal injection of ketamine (120 mg/kg; Vedco) and subcutaneous injection of xylazine (10.0 mg/kg; Akorn Inc). Animals were placed on a heating pad to maintain a constant body temperature at 37.5 ± 0.5 °C. The depth of anesthesia was confirmed by the absence of response to toe pinch. A laparotomy was performed, and the liver was gently moved dorsally to access the diaphragm. Gauze soaked with warm phosphate-buffered saline was placed on the liver to prevent dehydration. A handmade bipolar surface silver electrode was used to record spontaneous diaphragm EMG during normoxia (Li et al 2015b). EMGs were amplified (Model 1800; gain, 100; A-M Systems, Everett, WA, USA) and band pass-filtered (100 Hz to 10 kHz). The signals were digitized with an 8-channel Powerlab data acquisition device (Acquisition rate: 4 k/s; AD Instruments, Dunedin, New Zealand) connected to a computer and analyzed using LabChart 8 Pro software (AD Instruments, Dunedin, New Zealand; RRID:SCR_017551). The bilateral diaphragmatic EMGs were integrated twice (50 ms constant decay).

### Diaphragm whole-mount neuromuscular junction (NMJ) analysis

Following EMG recordings, diaphragm was dissected freshly and rinsed in phosphate buffered saline (PBS; pH 7.4) for NMJ analysis. The muscle was then fixed in 4% paraformaldehyde (PFA) in PBS at room temperature (RT) for 20 min. Connective tissue was removed, and diaphragms were incubated in 0.1 M glycine for 30 min at room temperature. Diaphragms were incubated with α-bungarotoxin Alexa Fluor 555 conjugate (1:200, Invitrogen) at RT for 15 min for postsynaptic nicotinic acetylcholine receptor labeling. Diaphragms were washed and permeabilized with ice-cold methanol for 5 min, and then blocked for 1h at RT in a Triton-bovine serum albumin (BSA)-PBS solution composed of 2% BSA and 0.2% Triton X-100 diluted in PBS. Diaphragms were incubated with presynaptic vesicle marker anti-SV2 antibody (1:10, Developmental Studies Hybridoma Bank, Iowa City, IA, USA; RRID:AB_2315387) and axonal marker anti-neurofilament antibody (1:1000, Clone: SMI-312, # 837904, BioLegend, San Diego, CA, USA; RRID:AB_2566782) at 4°C overnight. After washing with triton-BSA-PBS solution, diaphragms were incubated with Fluorescein (FITC) AffiniPure IgG, Fcγ subclass 1 specific secondary antibody (Jackson ImmunoResearch Laboratories, West Grove, PA, USA) for 1h at RT. Muscles were mounted with Vectashield mounting medium (Vector Laboratories), coverslipped and stored at −20°C. Whole-mount diaphragms were quantified using a Leica fluorescence microscope by two blinded individuals. Total numbers of NMJs and percentages of intact, partially-denervated, or completely-denervated NMJs were analyzed (Li et al 2015a). For each animal, 200–300 NMJs were quantified across the entire hemidiaphragm, and data were expressed for each NMJ phenotype as the percentage of total NMJs.

### Tissue processing

After diaphragm dissection, animals were intracardially perfused with heparinized saline (0.9% NaCl), followed by 4% PFA. Brain, brainstem and spinal cord were dissected out and postfixed at 4°C for 24h in 4% PFA. Tissues were then cryoprotected for 48 h in 30% sucrose (in 1X PBS).

### rVRG axon labeling in spinal cord

Frozen transverse spinal cord free-floating sections (40 µm) were cut using a Microm HM550 cryostat (Thermo Fisher Scientific, Waltham, MA, USA). Cutting order was maintained to distinguish slices from spinal cord rostral to lesion, at the lesion level, and caudal to lesion. Free-floating sections were then stored at 4°C in 0.1% azide in 1X PBS. Spinal cord sections were rinsed (3 x 5 min) in 1X PBS, then blocked for 30 min in blocking solution (5% normal donkey serum (NDS) and 0.2% triton in 1X PBS). They were then incubated for 4 days at 4°C in primary antibody solution (5% NDS and 0.1% triton in 1X PBS). The primary antibody used was rabbit-anti-dsRed (1:500, Living Colors DsRed pAb, RRID: AB_10013483; Takara Bio USA, Mountain View, CA, USA) to enhance mCherry-labeled rVRG axon visualization. Slices were washed 3 x 5 min and then incubated in secondary antibody solution (5% NDS and 0.1% triton in 1X PBS) for 1h at RT. The secondary antibody used was a donkey anti-rabbit Alexa Fluor 647 (1:200, A31573 Invitrogen, Carlsbad, CA, USA). Spinal cord was mounted on slides and coverslipped using mounting media. Images of the different sections (for each animal: rostral to lesion, at lesion level, and caudal to lesion) were captured with a Leica SP8 confocal microscope (Leica Microsystems Inc., Buffalo Grove, IL, USA) and LAS X imaging software (RRID: SCR_013673).

### rVRG axon quantification

Images were analyzed using ImageJ 1.53n software (NIH, USA; RRID: SCR_003070) and the SNT plugin (Arshadi et al 2021). Briefly, observed axons were delineated using SNT plugin, and axon profile numbers and total axon length were quantified (Cheng et al 2021a).

### Electrophysiology data processing

For EMGdia analysis, the amplitude of at least 10 double-integrated diaphragm EMG inspiratory bursts during normoxia was calculated from the injured side with LabChart 8 Pro software (AD Instruments, Colorado Springs, USA). We did not observe statistical differences between the animals anesthetized with isoflurane or ketamine/xylazine; therefore, we pooled the results obtained for each group and presented each animal value relative to the mean value for the DMSO-only group in the two anesthesia conditions (isoflurane and ketamine/xylazine). Diaphragm CMAPs for the injured side were averaged (at least 10 CMAP responses) and superimposed using LabChart 8 software (AD Instruments). The baseline-to-peak amplitude of the first wave of each superimposed CMAP was calculated.

### Experimental design and statistical analysis

All statistical analysis was performed using SigmaPlot 12.5 software (Systat Software, San Jose, CA, USA) (RRID: SCR_003210). One-way ANOVA was performed to compare the different groups for EMGdia, CMAP, NMJ and rVRG axon analyses. Statistical outliers were removed from the analyses using the interquartile range formula. All data were presented as mean ± S.E.M. Statistics were considered significant when *p* < 0.05. Before starting the study, rats were randomly assigned to experimental groups (n = 15-20 animals per group), and the different surgical procedures used within a given experiment were randomly distributed across these rats (and within a given surgical day). For the various analyses, we repeated the experiment in two large cohorts of animals. A timeline of the experimental design is shown in *Figure 1*. All surgical procedures and subsequent histological analyses were conducted in a blinded manner. In the Results section, we provide details of n’s, statistical tests used and the results of all statistical analyses (including exact p-values) for each experiment and for all statistical comparisons. In a separate Source Data file, we provide the raw data for each analysis in all the figures. We authenticated relevant experimental regents to ensure that they performed similarly across experiments and to validate the resulting data. For all antibodies used in the immunohistochemistry studies, we always verified (when receiving a new batch from the manufacturer) that labeling in the spinal cord and muscle coincided with the established expression pattern of the protein. We have provided Research Resource Identification Initiative (RRID) numbers for all relevant reagents (i.e. antibodies and computer programs) throughout the Materials and Methods section.

**Figure 1:**
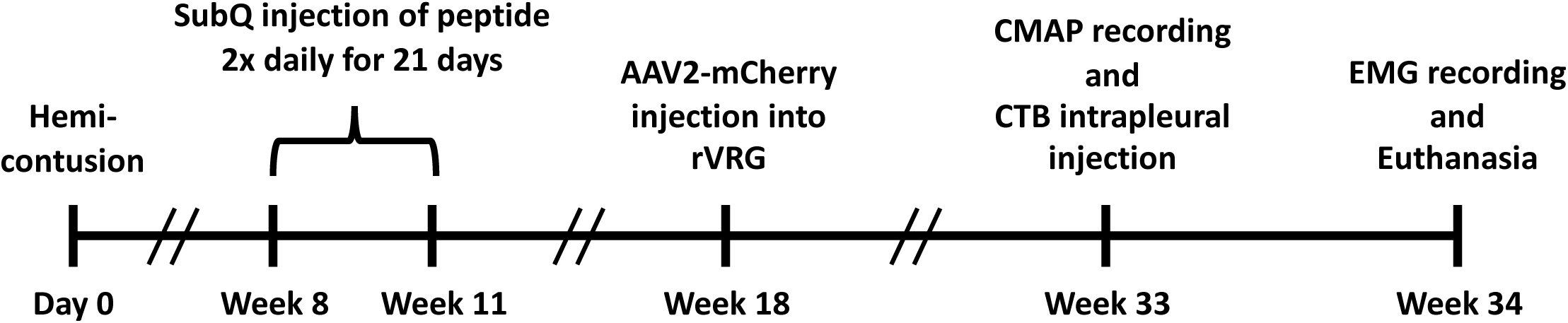
Experimental timeline. Rats received mid-cervical (C4/5) hemicontusion on day-0. Starting at 8 weeks post-SCI, rats were administered DMSO-only, PAP4, PTPσ peptide, or both peptides subcutaneously twice-daily for 21 days. At 33 weeks post-SCI, rats underwent intramedullary microinjection of AAV2-mCherry into ipsilesional rVRG. CMAP recordings were conducted on ipsilesional hemidiaphragm at week 36 post-injury, and CTB tracer was injected intrapleurally at this same time point. At 34 weeks after SCI, ipsilesional hemidiaphragm EMG recording was performed, followed by transcardial perfusion and tissue harvesting.

## RESULTS

### PAP4 promoted partial recovery of diaphragm motor activity in chronic cervical contusion SCI

To assess recovery of diaphragmatic respiratory function, we performed inspiratory electromyography (EMG) recording from diaphragm muscle in living animals during eupneic breathing. Previous work has shown that each of the three subregions of the hemidiaphragm is topographically innervated by PhMNs found at specific locations within cervical spinal cord: ventral subregion of the muscle is primarily innervated by PhMNs located at C3, medial subregion is innervated by C4 PhMNs, and dorsal subregion is controlled by PhMNs located at the C5 level (Laskowski & Sanes 1987). For this reason, we separately recorded EMGdia amplitude from ventral, medial and dorsal subregions of the ipsilesional hemidiaphragm at 34 weeks post-injury (P.I.) (n = 10-13 rats analyzed per group). Representative traces recorded from the dorsal portion of diaphragm are displayed for DMSO-only, PAP4, PTPσ peptide, and PAP4 + PTPσ peptide groups in *Figure 2A*. We did not observe any differences in EMGdia amplitude amongst the four conditions in ventral or medial subregions (p > 0.05 for all comparisons; *Figure 2B-C*). However, in the dorsal muscle subregion, we observed that the PAP4 group presented a significantly larger EMGdia amplitude compared to DMSO-only control animals (5.66 ± 6.08 A.U. vs 1.00 ± 1.03 A.U. respectively, p < 0.05; Kruskal-Wallis One Way ANOVA, Dunn’s Method; *Figure 2D*).

**Figure 2:**
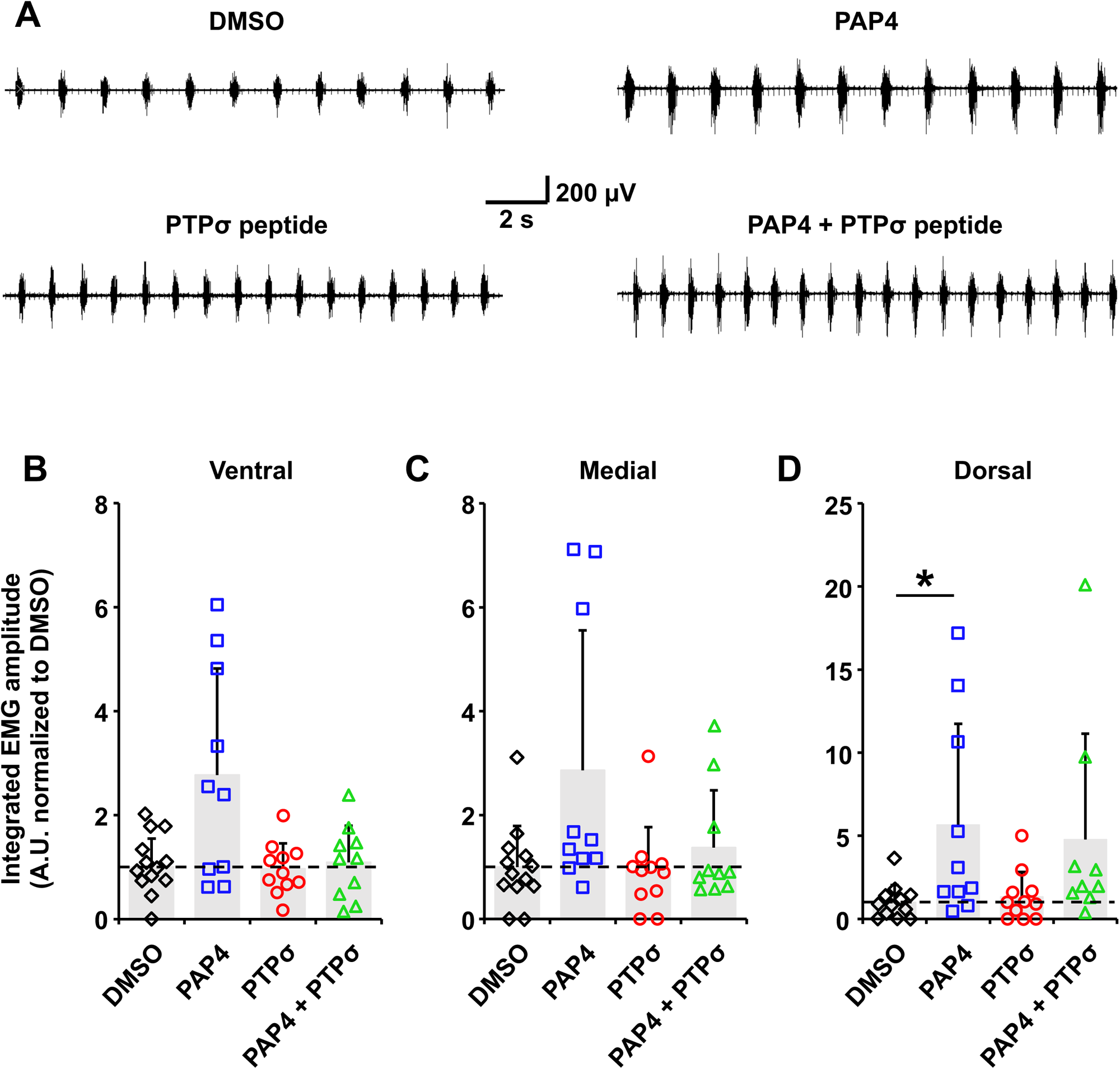
PAP4 promoted partial recovery of diaphragm EMG amplitude. (**A**) Representative traces of diaphragm electromyography (EMGdia) recordings ipsilateral to injury from dorsal diaphragm subregion for DMSO-only, PAP4, PTPσ peptide, and PAP4 + PTPσ peptide treated groups. Quantification of EMGdia amplitudes for ventral (**B**), medial (**C**) and dorsal (**D**) subregions of diaphragm ipsilateral to injury. * p < 0.05, Kruskal-Wallis One Way ANOVA (Dunn’s Method).

### PAP4 and PTPσ peptide did not impact functional diaphragm innervation

To examine whether peptide administration impacted diaphragm EMGdia recovery independently of effects on bulbospinal drive, we performed in vivo compound muscle action potential (CMAP) recordings from ipsilesional hemidiaphragm in response to supramaximal phrenic nerve stimulation to specifically assess functional diaphragm innervation by PhMNs. Unlike with EMGdia recordings that allow for individual analysis at each diaphragm subregion, CMAP recording simultaneously assesses the response across the entire hemidiaphragm. At 33 weeks P.I., we measured whole hemidiaphragm CMAP amplitude on the ipsilesional side (n = 11-17 rats analyzed per group). Representative traces are shown in *Figure 3A* for each condition. We did not observe any differences in CMAP amplitude amongst the different groups (p > 0.05 for all comparisons; Kruskal-Wallis One Way ANOVA, Dunn’s Method; *Figure 3B*), suggesting that the effect of PAP4 on functional EMGdia recovery was mediated by changes occurring centrally in the spinal cord and not in the peripheral portion of the rVRG-PhMN-diaphragm circuit.

**Figure 3:**
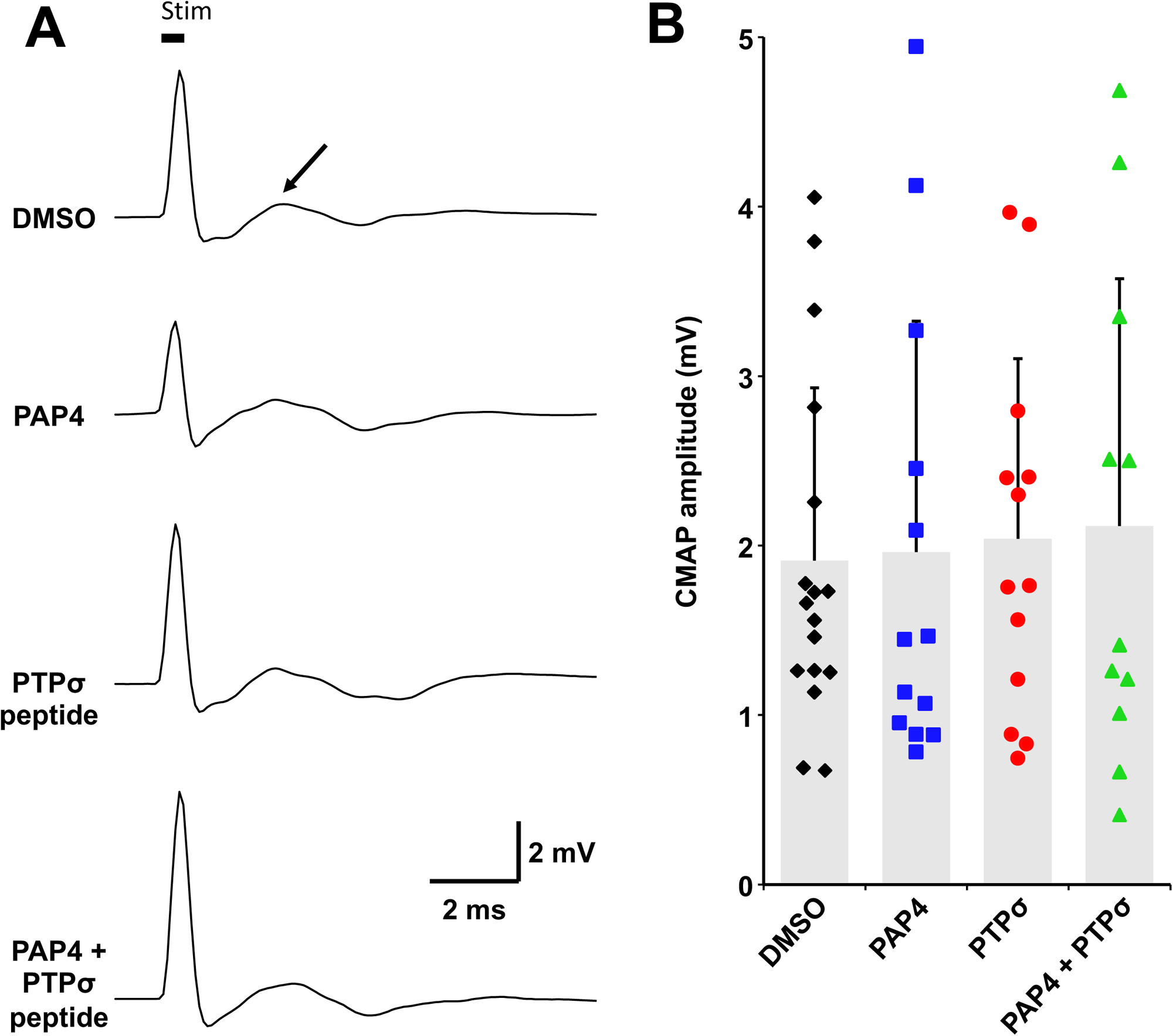
PAP4 and PTPσ peptide did not impact functional diaphragm innervation. (**A**) Representative diaphragm compound motor action potentials (CMAPs) recorded ipsilateral to injury for DMSO-only, PAP4, PTPσ peptide, and PAP4 + PTPσ peptide treated groups. Black arrow denotes a representative CMAP response. Black bar denotes the 0.5 ms stimulation of phrenic nerve. (**B**) Quantification of CMAP amplitudes for the different groups. p > 0.05 for all comparisons, Kruskal-Wallis One Way ANOVA (Dunn’s Method).

### PAP4 stimulated robust growth of bulbospinal rVRG respiratory axons within the phrenic nucleus

As we observed an effect of PAP4 treatment on diaphragmatic EMGs (which is a functional measure of rVRG-PhMN circuit connectivity) but not on diaphragm CMAPs (that instead assesses the connection from phrenic nerve to diaphragm), we then evaluated neuroanatomical changes centrally in the injured spinal cord that may be responsible for promoting functional improvement. Specifically, we examined the growth of ipsilateral-originating rVRG axons by microinjecting the anterograde tracer, AAV2-mCherry, into ipsilesional rVRG (*Figure 4A*). We performed this rVRG axon growth analysis at two different anatomical locations along the PhMN pool within the ipsilesional C3-C5 cervical spinal cord: (1) in the lesion site, and (2) in the intact C5 ventral horn caudal to the lesion (n = 5-6 rats analyzed per group). We observed that PAP4 promoted increased rVRG axon regrowth within the injury epicenter compared to DMSO-only control condition (*Figures 4C, D-G*) (quantification not shown). Importantly, we focused our rVRG axon analysis on the location within the phrenic nucleus caudal to the lesion, as these are the PhMNs that are significantly denervated from descending excitatory rVRG drive by the contusion (*Figure 4B, H-L*). We found a 5-8 fold increase in numbers of mCherry-labeled rVRG axon profiles in the PAP4-treated group (2.84 ± 1.25 per 100 µm^2^) compared to DMSO-only (0.46 ± 0.22, p = 0.002 vs PAP4) and PTPσ peptide (1.12 ± 1.34, p = 0.012 vs PAP4) groups, as well as in PAP4 + PTPσ peptide animals (2.89 ± 0.93) compared to DMSO-only (p = 0.02 vs combo) and PTPσ peptide (p = 0.013 vs combo) (Figure 4M). Similarly, the total length of mCherry^+^ rVRG axons was significantly higher in the PAP4-treated group (182.81 ± 100.50 µm per 100 µm^2^) compared to DMSO-only (22.93 ± 20.14, p = 0.002 vs PAP4) and PTPσ peptide (62.23 ± 80.91, p = 0.01 vs PAP4) groups, as well as in in PAP4 + PTPσ peptide animals (219.13 ± 26.20) compared to DMSO-only (p < 0.001 vs combo) and PTPσ peptide (p = 0.003 vs combo) (*Figure 4N*). As this cervical SCI model is a contusion that involves significant axon sparing on the side of injury, and is not an anatomically-complete hemisection, it is not known whether this rVRG axon growth at C5 is a product of regeneration, sprouting, or both.

**Figure 4:**
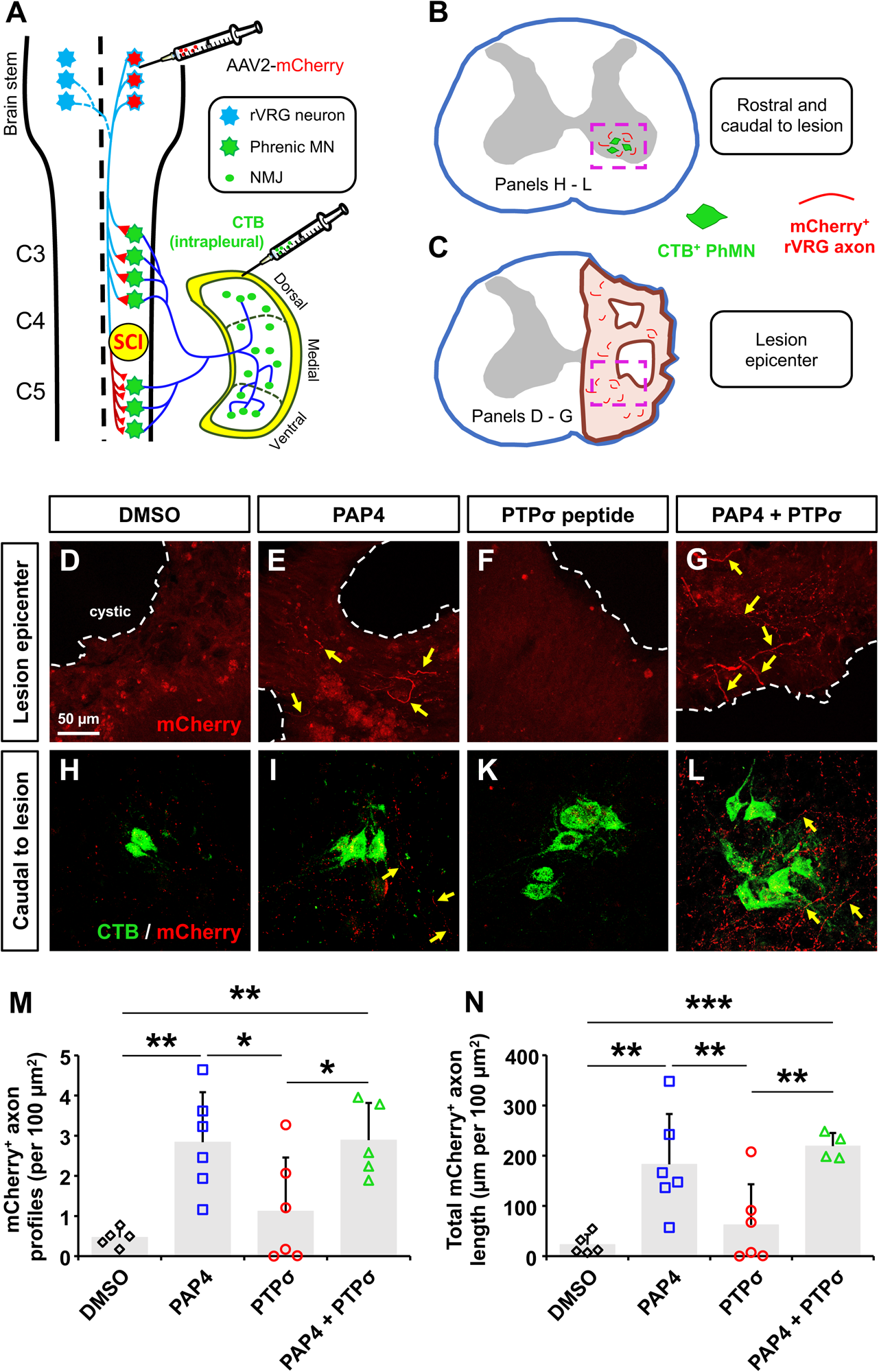
PAP4 promoted robust growth of rVRG axons within the phrenic nucleus. (**A**) AAV2-mCherry anterograde tracer was microinjected into ipsilesional rostral ventral respiratory group (rVRG). Labeled mCherry+ rVRG axons were observed surrounding CTB-labeled phrenic motoneurons located in C3-C5 spinal cord (**B**), as well as in the lesion site (**C**). Representative images are displayed for lesion epicenter (**D**-**G**) and ventral horn caudal to injury (**H**-**L**). Scale bar: 50µm (D-L). Quantification of mCherry+ axon profiles (**M**) and total length of mCherry+ axons (**N**) in the ipsilesional ventral horn caudal to injury at C5. * p ≤ 0.05, ** p ≤ 0.01, *** p ≤ 0.001, One Way ANOVA (Fisher LSD Method).

### PTPσ and PTEN inhibition impacted morphological innervation at the diaphragm NMJ

Mid-cervical contusion SCI disrupts rVRG-PhMN-diaphragm circuitry in two major ways: (1) the injury damages the rVRG axons, thereby removing control of PhMNs from excitatory rVRG drive (Urban et al 2018); and (2) the injury also leads to death of a portion of the PhMN cell bodies in cervical spinal cord, which results in partial denervation of diaphragm muscle (Nicaise et al 2012). While there is a major focus in the SCI field on trying to promote axon regrowth centrally in the injured spinal cord, inducing adaptive plasticity peripherally in the muscle is also highly important, though is not studied anywhere near as extensively. We therefore morphologically assessed peripheral nervous system effects of our peptide-based manipulations of PTEN and PTPσ on NMJ innervation by phrenic motor axons in ipsilesional hemidiaphragm (n = 8-14 rats analyzed per group). Representative NMJ images of postsynaptic nicotinic acetylcholine receptors labeled with α-bungarotoxin (red) and presynaptic motor axons labeled for both synaptic vesicle protein 2 and neurofilament (green) are shown in *Figure 5A-B*. High-magnification images of intact (*Figure 5C*), partially denervated (*Figure 5D*) and completely denervated (*Figure 5E*) NMJs are presented to show examples of quantified morphologies. In the ventral diaphragm subregion innervated by C3 PhMNs, no differences were observed amongst the four groups (p > 0.05 for all comparisons; Kruskal-Wallis One Way ANOVA, Dunn’s Method), with most of these NMJs remaining completely intact (*Figure 4F, I, L*). In the medial subregion, the percentage of intact NMJs was significantly lower for DMSO-only compared to PAP4 + PTPσ peptide (70.66 ± 21.25 % vs 90.91 ± 6.23 % respectively, p = 0.016), as well as for PTPσ peptide (56.66 ± 23.23 %) compared to both PAP4 (79.54 ± 15.48 %, p = 0.011 vs PTPσ) and PAP4 + PTPσ peptide (p < 0.001 vs PTPσ, One Way ANOVA (Fisher LSD Method); *Figure 5G*). In the medial portion, the PTPσ peptide group also presented significantly more partially denervated NMJs compared to PAP4 + PTPσ peptide (35.86 ± 21.04 % vs 7.02 ± 5.72 % respectively, p < 0.05, Kruskal-Wallis ANOVA (Dunn’s Method); *Figure 5J*), as well as more completely denervated NMJs compared to PAP4 + PTPσ peptide (8.44 ± 5.69 % vs 1.39 ± 2.09 % respectively, p < 0.05, Kruskal-Wallis ANOVA (Fisher LSD Method); *Figure 5M*). Finally, in the dorsal subregion, the percentage of intact NMJs was significantly lower for PTPσ peptide (36.63 ± 26.46 %) compared to PAP4 (72.97 ± 10.51 %, p < 0.05 vs PTPσ) and PAP4 + PTPσ peptide (82.99 ± 6.39 %, p < 0.05 vs PTPσ, Kruskal-Wallis ANOVA (Fisher LSD Method); *Figure 5H*). At this dorsal muscle location, the PTPσ peptide group also presented significantly more partially denervated NMJs compared to PAP4 + PTPσ peptide (51.40 ± 25.02 % vs 15,20 ± 10.29 % respectively, p < 0.05, Kruskal-Wallis ANOVA (Dunn’s Method); *Figure 5K*). No differences were observed amongst the four groups for totally denervated NMJs in the dorsal subregion (p > 0.05 for all comparisons; *Figure 5N*).

**Figure 5:**
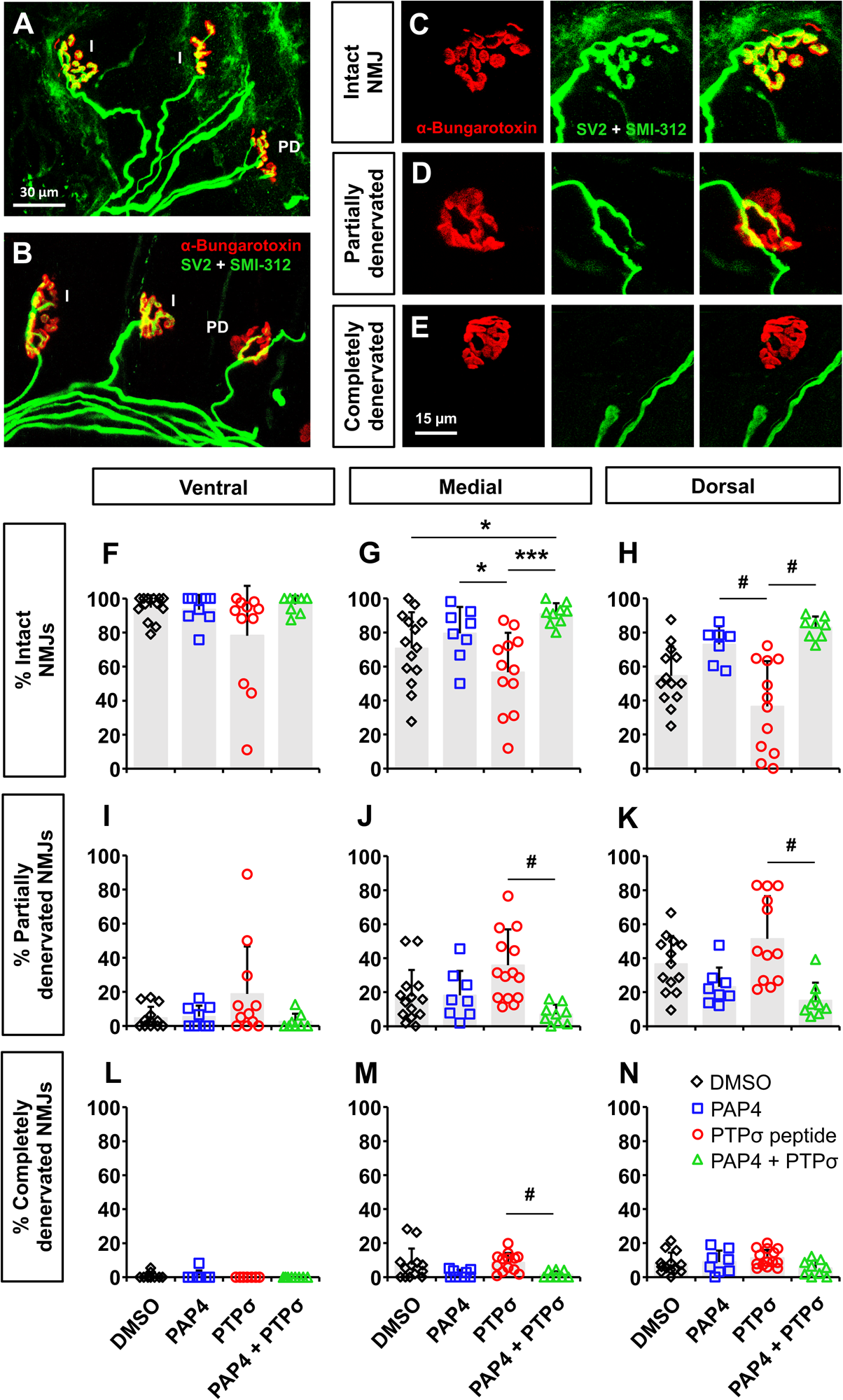
Morphological analysis of diaphragm neuromuscular junction (NMJ) innervation. Representative images illustrating examples of intact (I.) and partially denervated (P.D.) NMJs in ipsilesional hemidiaphragm (**A**-**B**), with enlargement images showing intact (**C**), partially denervated (**D**) and completely denervated (**E**) NMJs. Scale bar: 30µm (A-B), 15µm (C-E). Quantification of intact NMJs (**F**, **I**, **L**), partially denervated NMJs (**G**, **J**, **M**), and completely denervated NMJs (**H**, **K**, **N**) in ventral, medial and dorsal subregions. * p < 0.05, *** p < 0.001, One Way ANOVA (Fisher LSD Method); # p < 0.05, Kruskal-Wallis One Way ANOVA (Dunn’s Method).

### PTPσ inhibition induced the formation of atypical denervated diaphragm NMJ morphologies

Interestingly, we observed atypical morphologies of denervated NMJs with PTPσ peptide administration (*Figure 6A-B*) that we almost never find in models of muscle denervation, including contusion SCI. Specifically, in a large portion of NMJs in ipsilesional hemidiaphragm in only the PTPσ peptide condition, the phrenic motor axons regenerate to the synapse but appear to have an impaired ability to reform a synaptic connection with the previously-innervated muscle fiber. In these instances, thick SV2^+^/SMI-312+ pre-terminal phrenic motor axons overlay the sites of postsynaptic α-bungarotoxin^+^ nicotinic acetylcholine receptors, but fail to make a synaptic connection (*Figure 6A-B*). This atypical NMJ phenotype is virtually absent in the DMSO-only, PAP4-only and PAP4 + PTPσ peptide groups, but is found in a large percentage of NMJs in the PTPσ peptide-treated animals (PTPσ peptide: 13.73 ± 9.61 % of total NMJs; DMSO: 0.44 ± 0.76 %, p < 0.05 vs PTPσ peptide; PAP4: 0.59 ± 1.01 %, p < 0.05 vs PTPσ peptide; PAP4 + PTPσ peptide: 0.0 ± 0.0 %; p < 0.05 vs PTPσ peptide) (n = 5-8 rats analyzed per group) (*Figure 6C*).

**Figure 6:**
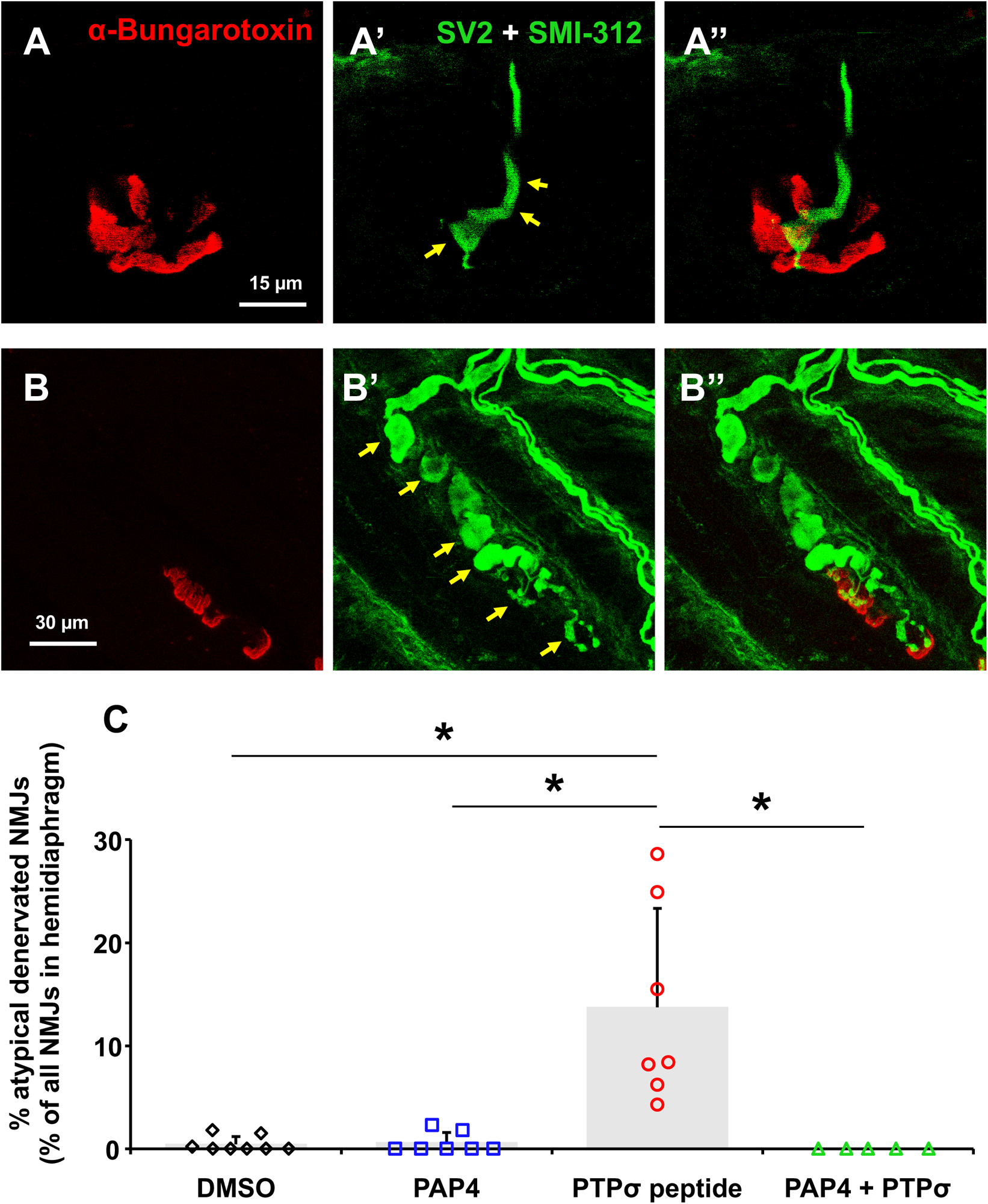
PTPσ peptide induced formation of atypical denervated neuromuscular junctions (NMJs). Representative atypical NMJ morphologies in ipsilesional hemidiaphragm of animals treated with PTPσ peptide (**A**-**B**). Scale bar: 15µm (A), 30µm (B). Yellow arrows denote thick SV2^+^/SMI-312^+^ pre-terminal phrenic motor axons (green) overlaying the sites of postsynaptic α-bungarotoxin^+^ nicotinic acetylcholine receptors but failing to make a synaptic connection (red). (**C**) Quantification of these atypical NMJs, expressed as percentage of total NMJs across the entire hemidiaphragm. * p < 0.05, Kruskal-Wallis One Way ANOVA (Dunn’s Method).

Collectively, these analyses of diaphragm NMJ innervation demonstrate that inhibition of PTPσ and PTEN (i.e. molecules which have been studied following nervous system injury primarily in the context of plasticity in the CNS) exert significant effects on axon populations at multiple locations in the damaged nervous system.

## DISCUSSION

In the present study, we tested the effects of systemic PAP4 and PTPσ inhibitory peptide administration on respiratory recovery following chronic cervical contusion SCI. We previously showed that these two drugs promote beneficial outcomes when delivered acutely/sub-acutely after cervical SCI (Urban et al 2020, Urban et al 2019), and we now report that PAP4 promotes significant axon growth and partial functional recovery in the context of chronic contusion SCI.

Rewiring of rVRG-PhMN-diaphragm circuitry is a highly important goal for achieving respiratory recovery following high SCI. This could be achieved through rVRG axon regeneration/sprouting and reconnection to PhMNs. However, there has been limited success in the identification of therapeutics to successfully promote diaphragm functional recovery after chronic SCI, and only a few studies to date have even focused on this objective (Cheng et al 2021a, Cheng et al 2021b, Tashiro et al 2017, Warren & Alilain 2018). Several approaches can be employed toward this goal. PTEN is a central inhibitor of neuron-intrinsic axon growth potential (Park et al 2010). Downregulation of PTEN activity following SCI has been shown to induce regrowth of adult axons using conditional PTEN knockout mice (Geoffroy et al 2015, Liu et al 2010, Metcalfe & Steward 2023, Park et al 2008, Sun et al 2011), shRNA vectors (Lewandowski & Steward 2014), and pharmacological inhibition (Ohtake et al 2014, Urban et al 2019). In the case of cervical SCI, we previously showed that pharmacologically inhibiting PTEN (using PAP4) at relatively early time points post-injury in a preclinical model of C2Hx leads to diaphragm activity recovery. This is associated with rVRG axons regenerating through the lesion and back toward their PhMN targets located in the C3 to C5 spinal cord (Urban et al 2019). More recently, we showed that this same treatment in the very same preclinical model, but delivered at a chronic stage following SCI, promotes minimal diaphragm activity recovery, though some rVRG axon growth with limited rVRG-PhMN synaptic reconnectivity could be observed (Cheng et al 2021a). Here, we used a model of mid-cervical hemicontusion, which contrary to the C2Hx model, leads to death of some PhMNs at the site of injury (while the C2Hx only results in denervation of these PhMNs from rVRG axon input) and also leaves some spared descending rVRG axons (while the C2Hx severs all ipsilesional rVRG axon input). The presence of these spared descending axons in the hemicontusion model used in the present study could explain why we observe a significant improvement in dorsal diaphragm activity associated with robustly increased rVRG axon growth caudal to injury (in the C5 spinal cord) with PAP4 peptide, but not previously in the C2Hx model (Cheng et al 2021a).

PTPσ plays a role in axon growth inhibition via direct binding to inhibitory molecules present in the injured spinal cord (Tran et al 2020). We previously showed that when delivered acutely following a C2Hx in rat, our PTPσ inhibitory peptide induces sprouting of spared contralateral-originating rVRG axons within the ipsilesional PhMN pool, in line with the observed diaphragm activity recovery (Urban et al 2020). Here, we used the same PTPσ peptide administration paradigm (i.e. same peptide concentration and similar delivery duration), but instead tested the effects in the chronic mid-cervical hemicontusion model. We do not observe any diaphragm activity recovery or enhanced rVRG axon growth in these rats treated with PTPσ peptide. The discrepancy between our past and present work could possibly be explained by the difference in timeline of administration. When peptide is delivered early after SCI (Urban et al 2019), axon growth inhibitory extracellular matrix, including CSPGs, is still in the process of increasing in density in the injured spinal cord, while it is already well-established in the present chronic protocol (Busch & Silver 2007), which further increases axon growth inhibition. We can hypothesize that, at this late time post-injury, the delivered PTPσ peptide is not effective in the face of these greater levels of neuronal-extrinsic inhibitory molecules (Milton et al 2023).

We also tested the effect of PAP4 + PTPσ peptide combination delivery, as we previously observed that they both induce partial diaphragm activity recovery when delivered individually at early time points post-injury (Urban et al 2020, Urban et al 2019). We hypothesized the effects would be amplified (either in an additive or even synergistic manner) compared to delivery of PAP4 alone or PTPσ peptide alone given that targeting PTEN and PTPσ in cervical SCI appears to exert effects on different modes of rVRG-PhMN circuit plasticity. We showed that subacute PAP4 delivery stimulates ipsilesional rVRG axon regeneration through injury to reach ipsilesional PhMNs located distal to the C2Hx (Urban et al 2019), while early PTPσ peptide delivery instead leads to robust sprouting of contralateral-originating rVRG axons and also of modulatory serotonergic axons within the ipsilesional PhMN pool distal to the C2Hx (Urban et al 2020). However, the diaphragm activity improvement in the combination condition is not significantly greater than PAP4 alone in the present study, and the combination effect on rVRG axon growth is also similar to PAP4-only treated animals. Considering these results, it is possible that the effects obtained with the two-peptide combination is due to solely to the PAP4. Though the aim of this study was to evaluate the effects of these molecules in the chronic injury, we can hypothesize that a potentiating effect may be observed if the PTPσ peptide is instead delivered at an early time point and then PAP4 is subsequently delivered at a chronic stage.

As contusion SCIs lead to focal lower motor neuron death, subsequent nerve degeneration and NMJ changes (Alvarez-Argote et al 2015, Nicaise et al 2013, Rana et al 2017), we evaluated whether our treatments have effects on NMJ innervation after cervical contusion SCI. Our results indicate that PAP4 + PTPσ peptide combination aids in preventing diaphragm denervation in our model. Surprisingly, PTPσ peptide alone significantly increases SCI-induced NMJ denervation. These changes in morphological diaphragm innervation are not reflected at the functional level using CMAP recording, which could be because CMAPs correspond to functional assessment across the entire hemidiaphragm in a single measurement, while NMJ morphologies can be assessed at a single-synapse level and specifically at different diaphragm subregions. In addition, we observed that PTPσ peptide treated animals present atypical NMJ morphologies (*Figure 6*), which have not been observed in the context of SCI to our knowledge. Furthermore, we are not aware of any previous SCI studies that morphologically assessed NMJ innervation following inhibition of PTEN or PTPσ. We can hypothesize that, since PTPσ is involved in motor axon (*i.e*., nerve) guidance during development (Sajnani-Perez et al 2003), inhibiting its function may lead to abnormal axon behavior at the NMJ after injury in the adult. The nature of these atypical NMJs and the reason they appear in response to PTPσ inhibition remain however to be further explored.

In this study, we report that the functional recovery achieved with PAP4 – while statistically significant in the dorsal subregion – is relatively modest. The degree of rVRG axon regrowth we observe with PTEN inhibition in this SCI model is greater than the level of functional recovery. One interpretation for this partial disconnect is that the circuitry requires more than just rVRG axon regrowth within the PhMN pool to produce functional improvement. As we previously showed in the chronic C2Hx model, PAP4 promoted robust rVRG axon regeneration, but little-to-no synaptic reinnervation of PhMNs located distal to the injury (Cheng et al 2021a). It is possible that a similar phenomenon is occurring in the present chronic contusion study where the growing rVRG axons are not synaptically reconnecting with their PhMN targets.

The results obtained in the present study illustrate how some treatments that lead to functional improvement when delivered at early time points post-SCI do not exert the same beneficial effects when delivered chronically. One option could be to adapt the treatment duration. Here, we employed the 21-day delivery paradigm we used previously that led to significant rVRG axon regeneration following C2Hx (Cheng et al 2021a). An extended period of treatment may lead to enhanced recovery in the chronic contusion model for PAP4 and/or PAP4 + PTPσ peptide.

A limitation of this work is the use of only female rats. We chose to use females because we have optimized the SCI model, PAP4 and PTPσ peptide treatments, and all of the axon track-tracing and functional assays in adult female rats in our previous studies (Charsar et al 2019, Cheng et al 2021a, Goulao et al 2019, Urban et al 2020, Urban et al 2019, Urban et al 2018). However, a recent study showed that male mice display greater recovery than females following SCI in response to PTEN deletion (Metcalfe & Steward 2023). It will be important to extend our work to male rats in future studies, especially considering that the majority of individuals impacted by SCI are men (Kumar et al 2018).

In conclusion, systemic administration of our PTEN-blocking peptide at a chronic stage following mid-cervical hemicontusion SCI promotes robust growth of descending bulbospinal respiratory axons that is associated with partial recovery of diaphragm function. Similar therapeutic results are observed with PAP4 + PTPσ peptide combination, while PTPσ inhibitory peptide alone exerted no beneficial effects. These exciting findings provide important information regarding the translation potential of PTEN-targeting therapeutics for repairing critically-important rVRG-PhMN-diaphragm circuitry in chronic cervical SCI.

## Acknowledgements

The authors would like to acknowledge Mr. Bob Yant for his incredible dedication to finding treatments that promote functional recovery for chronic spinal cord injury. His passion has been a major inspiration for this project, and this work was possible only through his generous support and constant encouragement.

## ABBREVIATIONS

AAV2: adeno-associated virus serotype 2
C2, C3, C4, C5, etc.: cervical level 2, 3, 4, 5, etc.
C2Hx: C2 spinal cord hemisection
ChABC: chondroitinase ABC
CNS: central nervous system
CMAP: compound motor action potential
CSPG: chondroitin sulfate proteoglycan
CTB: cholera toxin B subunit
EMG: electromyography
GAG: glycosaminoglycan
LAR: leukocyte common antigen-related phosphatase
mTOR: mammalian target of rapamycin
NMJ: neuromuscular junction
PAP4: PTEN antagonist peptide
PI3K: phosphatidylinositol 3-kinase
PhMN: phrenic motor neuron
PTEN: phosphatase and tensin homolog
PTPσ: protein tyrosine phosphatase sigma
rVRG: rostral ventral respiratory group
SCI: spinal cord injury

## REFERENCES

Ahuja CS, Wilson JR, Nori S, Kotter MRN, Druschel C, et al. 2017. Traumatic spinal cord injury. Nat. Rev. Dis. Primers 3: 17018

Alilain WJ, Horn KP, Hu H, Dick TE, Silver J. 2011. Functional regeneration of respiratory pathways after spinal cord injury. Nature 475: 196–200

Alvarez-Argote S, Gransee HM, Mora JC, Stowe JM, Jorgenson AJ, et al. 2015. The Impact of Midcervical Contusion Injury on Diaphragm Muscle Function. Journal of neurotrauma 33: 500–09

Arshadi C, Günther U, Eddison M, Harrington KIS, Ferreira TA. 2021. SNT: a unifying toolbox for quantification of neuronal anatomy. Nature Methods 18: 374–77

Bhowmick S, Abdul-Muneer PM. 2020. PTEN Blocking Stimulates Corticospinal and Raphespinal Axonal Regeneration and Promotes Functional Recovery After Spinal Cord Injury. Journal of Neuropathology & Experimental Neurology 80: 169–81

Bradbury EJ, Burnside ER. 2019. Moving beyond the glial scar for spinal cord repair. Nat Commun 10: 3879

Bradbury EJ, Moon LD, Popat RJ, King VR, Bennett GS, et al. 2002. Chondroitinase ABC promotes functional recovery after spinal cord injury. Nature 416: 636–40

Busch SA, Silver J. 2007. The role of extracellular matrix in CNS regeneration. Current Opinion in Neurobiology 17: 120–27

Charsar BA, Brinton MA, Locke K, Chen AY, Ghosh B, et al. 2019. AAV2-BDNF promotes respiratory axon plasticity and recovery of diaphragm function following spinal cord injury. The FASEB Journal 33: 13775–93

Cheng L, Sami A, Ghosh B, Goudsward HJ, Smith GM, et al. 2021a. Respiratory axon regeneration in the chronically injured spinal cord. Neurobiol Dis 155: 105389

Cheng L, Sami A, Ghosh B, Urban MW, Heinsinger NM, et al. 2021b. LAR inhibitory peptide promotes recovery of diaphragm function and multiple forms of respiratory neural circuit plasticity after cervical spinal cord injury. Neurobiol Dis 147: 105153

Cho YS, Kim SJ, Kim KH. 2020. Evaluation of PTEN Inhibitor Following Spinal Cord Injury on Recovery of Voiding Efficiency and Motor Function Observed by Regeneration in Spinal Cord. Int Neurourol J 24: 104–10

Danilov CA, Steward O. 2015. Conditional genetic deletion of PTEN after a spinal cord injury enhances regenerative growth of CST axons and motor function recovery in mice. Experimental neurology 266: 147–60

Dickendesher TL, Baldwin KT, Mironova YA, Koriyama Y, Raiker SJ, et al. 2012. NgR1 and NgR3 are receptors for chondroitin sulfate proteoglycans. Nature neuroscience 15: 703–12

Du K, Zheng S, Zhang Q, Li S, Gao X, et al. 2015. Pten Deletion Promotes Regrowth of Corticospinal Tract Axons 1 Year after Spinal Cord Injury. J Neurosci 35: 9754–63

Fisher D, Xing B, Dill J, Li H, Hoang HH, et al. 2011. Leukocyte Common Antigen-Related Phosphatase Is a Functional Receptor for Chondroitin Sulfate Proteoglycan Axon Growth Inhibitors. The Journal of Neuroscience 31: 14051–66

García-Alías G, Barkhuysen S, Buckle M, Fawcett JW. 2009. Chondroitinase ABC treatment opens a window of opportunity for task-specific rehabilitation. Nature neuroscience 12: 1145–51

Geoffroy CG, Lorenzana AO, Kwan JP, Lin K, Ghassemi O, et al. 2015. Effects of PTEN and Nogo Codeletion on Corticospinal Axon Sprouting and Regeneration in Mice. The Journal of Neuroscience 35: 6413–28

Giger RJ, Hollis ER, 2nd, Tuszynski MH. 2010. Guidance molecules in axon regeneration. Cold Spring Harb Perspect Biol 2: a001867

Goulao M, Ghosh B, Urban MW, Sahu M, Mercogliano C, et al. 2019. Astrocyte progenitor transplantation promotes regeneration of bulbospinal respiratory axons, recovery of diaphragm function, and a reduced macrophage response following cervical spinal cord injury. Glia 67: 452–66

Ito S, Ozaki T, Morozumi M, Imagama S, Kadomatsu K, Sakamoto K. 2021. Enoxaparin promotes functional recovery after spinal cord injury by antagonizing PTPRσ. Experimental neurology 340: 113679

James ND, Shea J, Muir EM, Verhaagen J, Schneider BL, Bradbury EJ. 2015. Chondroitinase gene therapy improves upper limb function following cervical contusion injury. Experimental neurology 271: 131–35

Kumar R, Lim J, Mekary RA, Rattani A, Dewan MC, et al. 2018. Traumatic Spinal Injury: Global Epidemiology and Worldwide Volume. World Neurosurg 113: e345–e63

Lang BT, Cregg JM, DePaul MA, Tran AP, Xu K, et al. 2015. Modulation of the proteoglycan receptor PTPσ promotes recovery after spinal cord injury. Nature 518: 404–8

Laskowski MB, Sanes JR. 1987. Topographic mapping of motor pools onto skeletal muscles. J Neurosci 7: 252–60

Lepore AC. 2011. Intraspinal cell transplantation for targeting cervical ventral horn in amyotrophic lateral sclerosis and traumatic spinal cord injury. J Vis Exp

Lepore AC, Tolmie C, O’Donnell J, Wright MC, Dejea C, et al. 2010. Peripheral hyperstimulation alters site of disease onset and course in SOD1 rats. Neurobiol Dis 39: 252–64

Lewandowski G, Steward O. 2014. AAVshRNA-mediated suppression of PTEN in adult rats in combination with salmon fibrin administration enables regenerative growth of corticospinal axons and enhances recovery of voluntary motor function after cervical spinal cord injury. J Neurosci 34: 9951–62

Li K, Hala TJ, Seetharam S, Poulsen DJ, Wright MC, Lepore AC. 2015a. GLT1 overexpression in SOD1(G93A) mouse cervical spinal cord does not preserve diaphragm function or extend disease. Neurobiol Dis 78: 12–23

Li K, Javed E, Hala TJ, Sannie D, Regan KA, et al. 2015b. Transplantation of glial progenitors that overexpress glutamate transporter GLT1 preserves diaphragm function following cervical SCI. Mol Ther 23: 533–48

Li K, Nicaise C, Sannie D, Hala TJ, Javed E, et al. 2014. Overexpression of the astrocyte glutamate transporter GLT1 exacerbates phrenic motor neuron degeneration, diaphragm compromise, and forelimb motor dysfunction following cervical contusion spinal cord injury. J Neurosci 34: 7622–38

Liu G, Detloff MR, Miller KN, Santi L, Houlé JD. 2012. Exercise modulates microRNAs that affect the PTEN/mTOR pathway in rats after spinal cord injury. Experimental neurology 233: 447–56

Liu K, Lu Y, Lee JK, Samara R, Willenberg R, et al. 2010. PTEN deletion enhances the regenerative ability of adult corticospinal neurons. Nat Neurosci 13: 1075–81

Luo X, Park KK. 2012. Neuron-intrinsic inhibitors of axon regeneration: PTEN and SOCS3. Int Rev Neurobiol 105: 141–73

Martin M, Li K, Wright MC, Lepore AC. 2015. Functional and morphological assessment of diaphragm innervation by phrenic motor neurons. J Vis Exp: e52605

Metcalfe M, Steward O. 2023. PTEN deletion in spinal pathways via retrograde transduction with AAV-RG enhances forelimb motor recovery after cervical spinal cord injury; Sex differences and late-onset pathophysiologies. Experimental neurology 370: 114551

Milton AJ, Kwok JCF, McClellan J, Randall SG, Lathia JD, et al. 2023. Recovery of Forearm and Fine Digit Function After Chronic Spinal Cord Injury by Simultaneous Blockade of Inhibitory Matrix Chondroitin Sulfate Proteoglycan Production and the Receptor PTPσ. Journal of neurotrauma 40: 2500–21

Nicaise C, Frank DM, Hala TJ, Authelet M, Pochet R, et al. 2013. Early phrenic motor neuron loss and transient respiratory abnormalities after unilateral cervical spinal cord contusion. J Neurotrauma 30: 1092–9

Nicaise C, Hala TJ, Frank DM, Parker JL, Authelet M, et al. 2012. Phrenic motor neuron degeneration compromises phrenic axonal circuitry and diaphragm activity in a unilateral cervical contusion model of spinal cord injury. Experimental neurology 235: 539–52

Ohtake Y, Park D, Abdul-Muneer PM, Li H, Xu B, et al. 2014. The effect of systemic PTEN antagonist peptides on axon growth and functional recovery after spinal cord injury. Biomaterials 35: 4610–26

Park KK, Liu K, Hu Y, Kanter JL, He Z. 2010. PTEN/mTOR and axon regeneration. Experimental neurology 223: 45–50

Park KK, Liu K, Hu Y, Smith PD, Wang C, et al. 2008. Promoting axon regeneration in the adult CNS by modulation of the PTEN/mTOR pathway. Science 322: 963–6

Ramer LM, Ramer MS, Bradbury EJ. 2014. Restoring function after spinal cord injury: towards clinical translation of experimental strategies. The Lancet Neurology 13: 1241–56

Rana S, Sieck GC, Mantilla CB. 2017. Diaphragm electromyographic activity following unilateral midcervical contusion injury in rats. J Neurophysiol 117: 545–55

Rosenzweig ES, Salegio EA, Liang JJ, Weber JL, Weinholtz CA, et al. 2019. Chondroitinase improves anatomical and functional outcomes after primate spinal cord injury. Nature neuroscience 22: 1269–75

Rowland JW, Hawryluk GW, Kwon B, Fehlings MG. 2008. Current status of acute spinal cord injury pathophysiology and emerging therapies: promise on the horizon. Neurosurg Focus 25: E2

Sajnani-Perez G, Chilton JK, Aricescu AR, Haj F, Stoker AW. 2003. Isoform-specific binding of the tyrosine phosphatase ptpσ to a ligand in developing muscle. Molecular and Cellular Neuroscience 22: 37–48

Shen Y, Tenney AP, Busch SA, Horn KP, Cuascut FX, et al. 2009. PTPsigma is a receptor for chondroitin sulfate proteoglycan, an inhibitor of neural regeneration. Science 326: 592–6

Stewart AN, Kumari R, Bailey WM, Glaser EP, Hammers GV, et al. 2023. PTEN knockout using retrogradely transported AAVs restores locomotor abilities in both acute and chronic spinal cord injury. bioRxiv: 2023.04.17.537179

Sun F, Park KK, Belin S, Wang D, Lu T, et al. 2011. Sustained axon regeneration induced by co-deletion of PTEN and SOCS3. Nature 480: 372–5

Tashiro S, Nakamura M, Okano H. 2017. The prospects of regenerative medicine combined with rehabilitative approaches for chronic spinal cord injury animal models. Neural Regen Res 12

Tester NJ, Howland DR. 2008. Chondroitinase ABC improves basic and skilled locomotion in spinal cord injured cats. Experimental neurology 209: 483–96

Thuret S, Moon LDF, Gage FH. 2006. Therapeutic interventions after spinal cord injury. Nature Reviews Neuroscience 7: 628–43

Tonks NK. 2006. Protein tyrosine phosphatases: from genes, to function, to disease. Nat Rev Mol Cell Biol 7: 833–46

Tran AP, Warren PM, Silver J. 2020. Regulation of autophagy by inhibitory CSPG interactions with receptor PTPσ and its impact on plasticity and regeneration after spinal cord injury. Experimental neurology 328: 113276

Urban MW, Ghosh B, Block CG, Charsar BA, Smith GM, et al. 2020. Protein Tyrosine Phosphatase sigma Inhibitory Peptide Promotes Recovery of Diaphragm Function and Sprouting of Bulbospinal Respiratory Axons after Cervical Spinal Cord Injury. J Neurotrauma

Urban MW, Ghosh B, Block CG, Strojny LR, Charsar BA, et al. 2019. Long-Distance Axon Regeneration Promotes Recovery of Diaphragmatic Respiratory Function after Spinal Cord Injury. eNeuro 6

Urban MW, Ghosh B, Strojny LR, Block CG, Blazejewski SM, et al. 2018. Cell-type specific expression of constitutively-active Rheb promotes regeneration of bulbospinal respiratory axons following cervical SCI. Exp Neurol 303: 108–19

Venkatesh K, Ghosh SK, Mullick M, Manivasagam G, Sen D. 2019. Spinal cord injury: pathophysiology, treatment strategies, associated challenges, and future implications. Cell and Tissue Research 377: 125–51

Walker CL, Xu X-M. 2014. PTEN inhibitor bisperoxovanadium protects oligodendrocytes and myelin and prevents neuronal atrophy in adult rats following cervical hemicontusive spinal cord injury. Neuroscience Letters 573: 64–68

Warren PM, Alilain WJ. 2018. Plasticity Induced Recovery of Breathing Occurs at Chronic Stages after Cervical Contusion. Journal of neurotrauma 36: 1985–99

Warren PM, Steiger SC, Dick TE, MacFarlane PM, Alilain WJ, Silver J. 2018. Rapid and robust restoration of breathing long after spinal cord injury. Nat. Commun. 9: 4843–43

Winslow C, Rozovsky J. 2003. Effect of spinal cord injury on the respiratory system. Am J Phys Med Rehabil 82: 803–14

Xie X, Luo L, Liang M, Zhang W, Zhang T, et al. 2020. Structural basis of liprin-α-promoted LAR-RPTP clustering for modulation of phosphatase activity. Nat Commun 11: 169

Xu B, Park D, Ohtake Y, Li H, Hayat U, et al. 2015. Role of CSPG receptor LAR phosphatase in restricting axon regeneration after CNS injury. Neurobiology of disease 73: 36–48

Zhu H, Xie R, Liu X, Shou J, Gu W, et al. 2017. MicroRNA-494 improves functional recovery and inhibits apoptosis by modulating PTEN/AKT/mTOR pathway in rats after spinal cord injury. Biomedicine & Pharmacotherapy 92: 879–87

Zukor K, Belin S, Wang C, Keelan N, Wang X, He Z. 2013. Short hairpin RNA against PTEN enhances regenerative growth of corticospinal tract axons after spinal cord injury. J Neurosci 33: 15350–61

